# Biogeographic dating of speciation times using paleogeographically informed processes

**DOI:** 10.1101/028738

**Authors:** Michael J. Landis

## Abstract

Standard models of molecular evolution cannot estimate absolute speciation times alone, and require external calibrations to do so. Because fossil calibration methods rely on the unreliable fossil record, most nodes in the tree of life are dated with poor accuracy. However, many major paleogeographical events are dated, and since biogeographic processes depend on paleogeographical conditions, biogeographic dating may be used as an alternative or complementary method to fossil dating. I demonstrate how a time-stratified biogeographic stochastic process may be used to estimate absolute divergence times by conditioning on dated paleogeographical events. Informed by the current paleogeographical literature, I construct an empirical dispersal graph using 25 areas and 26 epochs for the past 540 Ma of Earth’s history. Simulations indicate biogeographic dating performs well so long as paleogeography imposes constraint on biogeographic character evolution. To gauge whether biogeographic dating may have any practical use, I analyze the well-studied turtle clade (*Testudines*) then assess how well biogeographic dating fares compared to heavily fossil-calibrated dating results as reported in the literature. Fossil-free biogeographic dating estimated the age of the most recent common ancestor of extant turtles to be approximately 201 Ma, which is consistent with fossil-based estimates. Accuracy improves further when including a root node fossil calibration. The described model, paleogeographical dispersal graph, and analysis scripts are available for use with RevBayes.

## 1 Introduction

Time is a simple and fundamental axis of evolution. Knowing the order and timing of evolutionary events grants us insight into how vying evolutionary processes interact. With a perfectly accurate catalog of geologically-dated speciation times, many macroevolutionary questions would yield to simple interrogation, such as whether one clade exploded with diversity before or after a niche-analogous clade went extinct, or whether some number of contemporaneous biota were eradicated simultaneously by the same mass extinction event. Only rarely does the fossil record give audience to the exact history of evolutionary events: it is infamously irregular across time, space, and species, so biologists generally resort to inference to estimate when, where, and what happened to fill those gaps. That said, we have not yet found a perfect character or model to infer dates for divergence times, so advances in dating strategies are urgently needed. A brief survey of the field reveals why.

The molecular clock hypothesis of Zuckerkandl and Pauling (1962) states that if substitutions arise (i.e. alleles fix) at a constant rate, the expected number of substitutions is the product of the substitution rate and the time the substitution process has been operating. With data from extant taxa, we only observe the outcome of the evolutionary process for an unknown rate and an unknown amount of time. As such, rate and time are non-identifiable under standard models of molecular substitution, so inferred amounts of evolutionary change are often reported as a compound parameter, the product of rate and time, called length. If all species’ shared the same substitution rate, a phylogeny with branches measured in lengths would give relative divergence times, i.e. proportional to absolute divergence times. While it is reasonable to say species’ evolution shares a basis in time, substitution rates differ between species and over macroevolutionary timescales (Wolfe et al. 1987; Martin and Palumbi 1993). Even when imposing a model of rate heterogeneity (Thorne et al. 1998; Drummond et al. 2006), only relative times may be estimated. Extrinsic information, i.e. a dated calibration point, is needed to establish an absolute time scaling, and typically takes form as a fossil occurrence or paleogeographical event.

When fossils are available, they currently provide the most accurate inroad to calibrate divergence events to geological timescales, largely because each fossil is associated with a geological occurrence time. Fossil ages may be used in several ways to calibrate divergence times. The simplest method is the fossil node calibration, whereby the fossil is associated with a clade and constrains its time of origin (Ho and Phillips 2009; Parham et al. 2011). Node calibrations are empirical priors, not data-dependent stochastic processes, so they depend entirely on experts’ abilities to quantify the distribution of plausible ages for the given node. That is, node calibrations do not arise from a generative evolutionary process. Since molecular phylogenies cannot identify rate from time, then the time scaling is entirely determined by the prior, i.e. the posterior is perfectly prior-sensitive for rates and times. Rather than using prior node calibrations, fossil tip dating (Pyron 2011; Ronquist et al. 2012) treats fossil occurrences as terminal taxa with morphological characters as part of any standard phylogenetic analysis. In this case, the model of character evolution, tree prior, and fossil ages generate the distribution of clade ages, relying on the fossil ages and a morphological clock to induce time calibrations. To provide a generative process of fossilization, Heath et al. (2014) introduced the fossilized birth-death process, by which lineages speciate, go extinct, or produce fossil observations. Using fossil tip dating with the fossilized birth-death process, Gavryushkina et al. (2015) demonstrated multiple calibration techniques may be used jointly in a theoretically consistent framework (i.e. without introducing model violation).

Of course, fossil calibrations require fossils, but many clades leave few to no known fossils due to taphonomic processes, which filter out species with too soft or too fragile of tissues, or with tissues that were buried in substrates that were too humid, too arid, or too rocky; or due to sampling biases, such as geographical or political biases imbalancing collection efforts (Behrensmeyer et al. 2000; Kidwell and Holland 2002). Although these biases do not prohibitively obscure the record for widespread species with robust mineralized skeletons—namely, large vertebrates and marine invertebrates—fossil-free calibration methods are desperately needed to date the remaining majority of nodes in the tree of life.

In this direction, analogous to fossil node dating, node dates may be calibrated using paleobiogeographic scenarios (Heads 2005; Renner 2005). For example, an ornithologist might reasonably argue that a bird known as endemic to a young island may have speciated only after the island was created, thus providing a maximum age of origination. Using this scenario as a calibration point excludes the possibility of alternative historical biogeographic explanations, e.g. the bird might have speciated off-island before the island surfaced and migrated there afterwards. See Heads (2005; 2011), Kodandaramaiah (2011), and Ho et al. (2015) for discussion on the uses and pitfalls of biogeographic node calibrations. Like fossil node calibrations, biogeographic node calibrations fundamentally rely on some prior distribution of divergence times, opinions may vary from expert to expert, making results difficult to compare from a modeling perspective. Worsening matters, the time and context of biogeographic events are never directly observed, so asserting that a particular dispersal event into an island system resulted in a speciation event to calibrate a node fails to account for the uncertainty that the assumed evolutionary scenario took place at all. Ideally, all possible biogeographic and diversification scenarios would be considered, with each scenario given credence in proportion to its probability.

Inspired by advents in fossil dating models (Pyron 2011; Ronquist et al. 2012; Heath et al. 2014), which have matured from phenomenological towards mechanistic approaches (Rodrigue and Philippe 2010), I present an explicitly data-dependent and process-based biogeographic method for divergence time dating to formalize the intuition underlying biogeographic node calibrations. Analogous to fossil tip dating, the goal is to allow the observed biogeographic states at the “tips” of the tree induce a posterior distribution of dated speciation times by way of an evolutionary process. By modeling dispersal rates between areas as subject to time-calibrated paleogeographical information, such as the merging and splitting of continental adjacencies due to tectonic drift, particular dispersal events between area-pairs are expected to occur with higher probability during certain geological time intervals than during others. For example, the dispersal rate between South America and Africa was likely higher when they were joined as West Gondwana (ca 120 Ma) than when separated as they are today. If the absolute timing of dispersal events on a phylogeny matters, then so must the absolute timing of divergence events. Unlike fossil tip dating, biogeographic dating should, in principle, be able to date speciation times only using extant taxa.

To illustrate how this is possible, I construct a toy biogeographic example to demonstrate when paleogeography may date divergence times, then follow with a more formal description of the model. By performing joint inference with molecular and biogeographic data, I demonstrate the effectiveness of biogeographic dating by applying it to simulated and empirical scenarios, showing rate and time are identifiable. While researchers have accounted for phylogenetic uncertainty in biogeographic analyses (Nylander et al. 2008; Lemey et al. 2009; Beaulieu et al. 2013), I am unaware of work demonstrating how paleogeographic calibrations may be leveraged to date divergence times via a biogeographic process. For the empirical analysis, I date the divergence times for *Testudines* using biogeographic dating, first without any fossils, then using a fossil root node calibration. Finally, I discuss the strengths and weaknesses of my method, and how it may be improved in future work.

## 2 Model

### 2.1 The anatomy of biogeographic dating

Briefly, I will introduce an example of how time-calibrated paleogeographical events may impart information through a biogeographic process to date speciation times, then later develop the details underlying the strategy, which I call biogeographic dating. Throughout the manuscript, I assume a rooted phylogeny, Ψ, with known topology but with unknown divergence times that I wish to estimate. Time is measured in geological units and as time until present, with *t* = 0 being the present, *t* < 0 being the past, and age being the negative amount of time until present. To keep the model of biogeographic evolution simple, the observed taxon occurrence matrix, **Z**, is assumed to be generated by a discrete-valued dispersal process where each taxon is present in only a single area at a time (Sanmartín et al. 2008). For example, taxon *T*1 might be coded to be found in Area *A* or Area *B* but not both simultaneously. Although basic, this model is sufficient to make use of paleogeographical information, suggesting more realistic models will fare better.

Consider two areas, *A* and *B*, that drift into and out of contact over time. When in contact, dispersal is possible; when not, impossible. Represented as a graph, *A* and *B* are vertices, and the edge (*A, B*) exists only during time intervals when *A* and *B* are in contact. The addition and removal of dispersal routes demarcate time intervals, or *epochs*, each corresponding to some epoch index, *k* ∈ {1,…, *K*}. To define how dispersal rates vary with *k*, I use an epoch-valued continuous-time Markov chain (CTMC) (Ree et al. 2005; Ree and Smith 2008; Bielejec et al. 2014). The adjacency matrix for the *k*^th^ time interval’s graph is used to populate the elements of an instantaneous rate matrix for an epoch’s CTMC such that the dispersal rate is equal to 1 when the indexed areas are adjacent and equals 0 otherwise. For a time-homogeneous CTMC, the transition probability matrix is typically written as **P**(*t*), which assumes the rate matrix, **Q**, has been rescaled by some clock rate, *μ*, and applied to a branch of some length, *t*. For a time-heterogeneous CTMC, the value of the rate matrix changes as a function of the underlying time interval, **Q**(*k*). The transition probability matrix for the time-heterogeneous process, **P**(*s*,*t*), is the matrix-product of the constituent epochs’ time-homogeneous transition probability matrices, and takes a value determined by the absolute time and order of paleogeographical events contained between the start time, *s*, and end time, *t*. Under this construction, certain types of dispersal events are more likely to occur during certain absolute (not relative) time intervals, which potentially influences probabilities of divergence times in absolute units.

Below, I give examples of when a key divergence time is likely to precede a split event (Figure 1) or to follow a merge event (Figure 2). To simplify the argument, I assume a single change must occur on a certain branch given the topology and tip states, though the logic holds in general.

**Figure 1:**
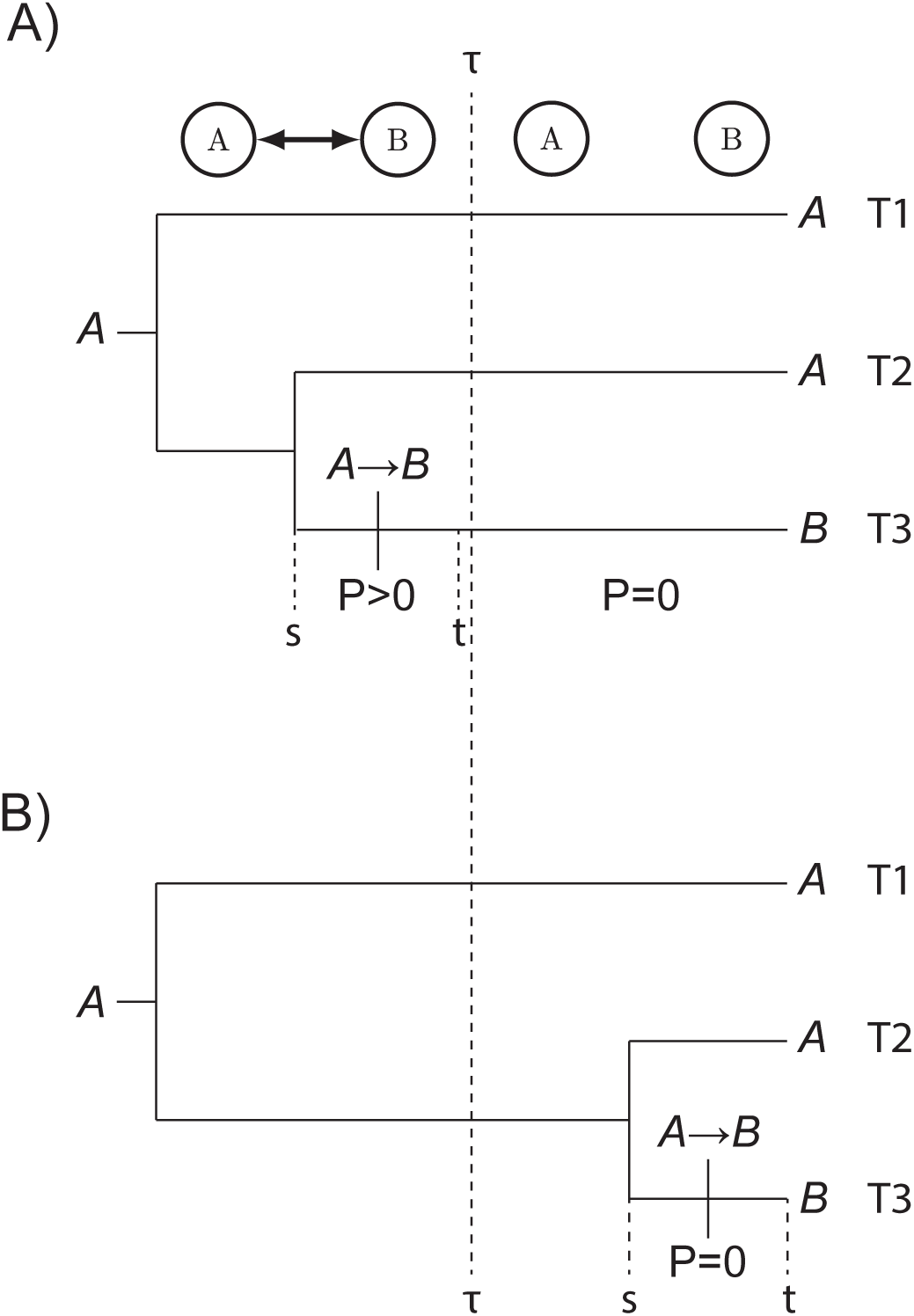
Effects of a paleogeographical split on divergence times. Area *A* splits from Area *B* at time *τ. T*1 and *T*2 have state *A* and the transition *A* → *B* most parsimoniously explains how *T*3 has state *B*. The transition probabilty for *P* = [**P**(*s*, *t*)] ab is non-zero before the paleogeographical split event at time *τ*, and is zero afterwards. Two possible divergence and dispersal times are given: A) *T*3 originates before the split when the transition *A* → *B* has non-zero probability. B) *T*3 originates after the split when the transition *A* → *B* has probability zero.

In the first scenario (Figure 1), sister taxa *T*2 and *T*3 are found in Areas *A* and *B*, respectively. The divergence time, *s*, is a random variable to be inferred. At time *τ*, the dispersal route (*A, B*) is destroyed, inducing the transition probability [**P**(*s, t*)]*_AB_* = 0 between times *τ* and 0. Since *T*2 and *T*3 are found in different areas, at least one dispersal event must have occurred during an interval of non-zero dispersal probability. Then, the divergence event that gave rise to *T*2 an *T*3 must have also pre-dated *τ*, with at least one dispersal event occuring before the split event (Figure 1A). If *T*2 and *T*3 diverge after *τ*, a dispersal event from *A* to *B* is necessary to explain the observations (Figure 1B), but the model disfavors that divergence time because the required transition has probability zero. In this case, the creation of a dispersal barrier informs the latest possible divergence time, a bound after which divergence between *T*2 and *T*3 is distinctly less probable if not impossible. It is also worth considering that a more complex process modeling vicariant speciation would provide tight bounds centered on *τ* (see Discussion).

In the second scenario (Figure 2), the removal of a dispersal barrier is capable of creating a maximum divergence time threshold, pushing divergence times towards the present. To demonstrate this, say the ingroup sister taxa *T*3 and *T*4 both inhabit Area *B* and the root state is Area *A*. Before the areas merge, the rate of dispersal between *A* and *B* as zero, and non-zero afterwards. When speciation happens after the areas merge, then the ancestor of (*T*3, *T*4) may disperse from *A* to *B*, allowing *T*3 and *T*4 to inherit state *B* (Figure 2A). Alternatively, if *T*3 and *T*4 originate before the areas merge, then the same dispersal event on the branch ancestral to (T3,T4) has probability zero (Figure 2B).

**Figure 2:**
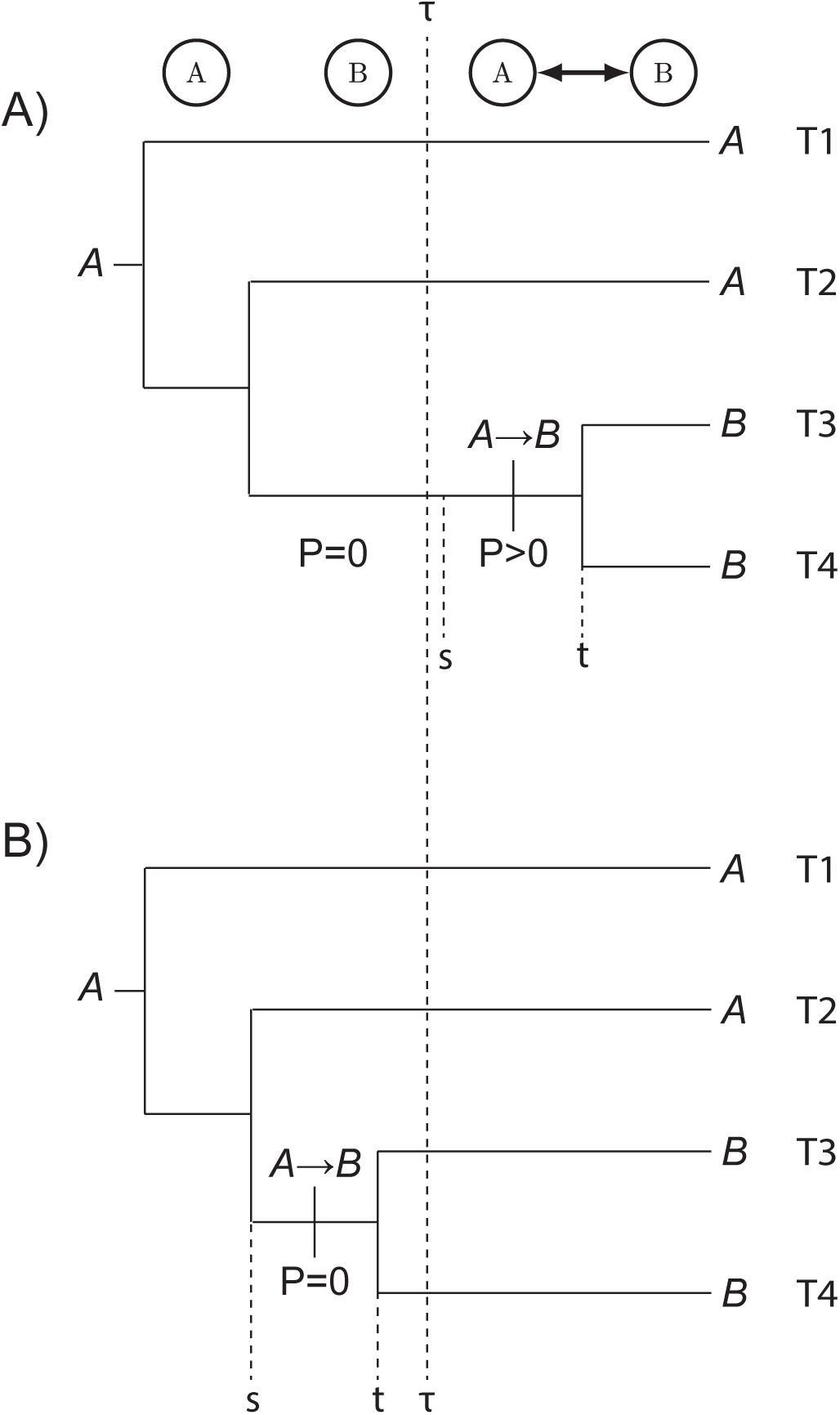
Effects of paleogeographical merge on divergence times. Area *A* merges with Area *B* at time *τ*. *T*1 and *T*2 have the state *A* and the transition *A* → *B* on the lineage leading to (*T*3, *T*4) most parsimoniously explains how *T*3 and *T*4 have state *B*. The transition probabilty for *P* = [**P**(*s*, *t*)]*_AB_* is zero before the paleogeographical merge event at time *τ*, and only non-zero afterwards. Two possible divergence and dispersal times are given: A) *T*3 and *T*4 originate after the merge when the transition *A* → *B* has non-zero probability. B) *T*3 and *T*4 originate before the merge when the transition *A* → *B* has probability zero.

### 2.2 Paleogeography, graphs, and Markov chains

How biogeography may date speciation times depends critically on the assumptions of the biogeographic model. The above examples depend on the notion of *reachability*, that two vertices (areas) are connected by some ordered set of edges (dispersal routes) of any length. In the adjacent-area dispersal model used here, one area might not be reachable from another area during some time interval, during which the corresponding transition probability is zero. That is, no path of any number of edges (series of dispersal events) may be constructed to connect the two areas. The concept of reachability may be extended to sets of partitioned areas: in graph theory, sets of vertices (areas) that are mutually reachable are called *(connected) components*. In terms of a graphically structured continuous time Markov chain, each component forms a *communicating class*: a set of states with positive transition probabilities only to other states in the set, and zero transition probabilities to other states (or communicating classes) in the state space. To avoid confusion with the “generic” biogeographical concept of components (Passalacqua 2015), and to emphasize the interaction of these partitioned states with respect to the underlying stochastic process, I hereafter refer to these sets of areas as communicating classes.

Taking terrestrial biogeography as an example, areas exclusive to Gondwana or Laurasia may each reasonably form communicating classes upon the break-up of Pangaea (Figure 3), meaning species are free to disperse within these paleocontinents, but not between them. For example, the set of communication classses is *S* = {{Afr}, {As}, {Ind}} at *t* = −100, i.e. there are |*S*| = 3 communicating classes because no areas share edges (Figure 3C), while at *t* = −10 there is |*S*| = 1 communicating class since a path exists between all three pairs of areas (Figure 3E).

**Figure 3:**
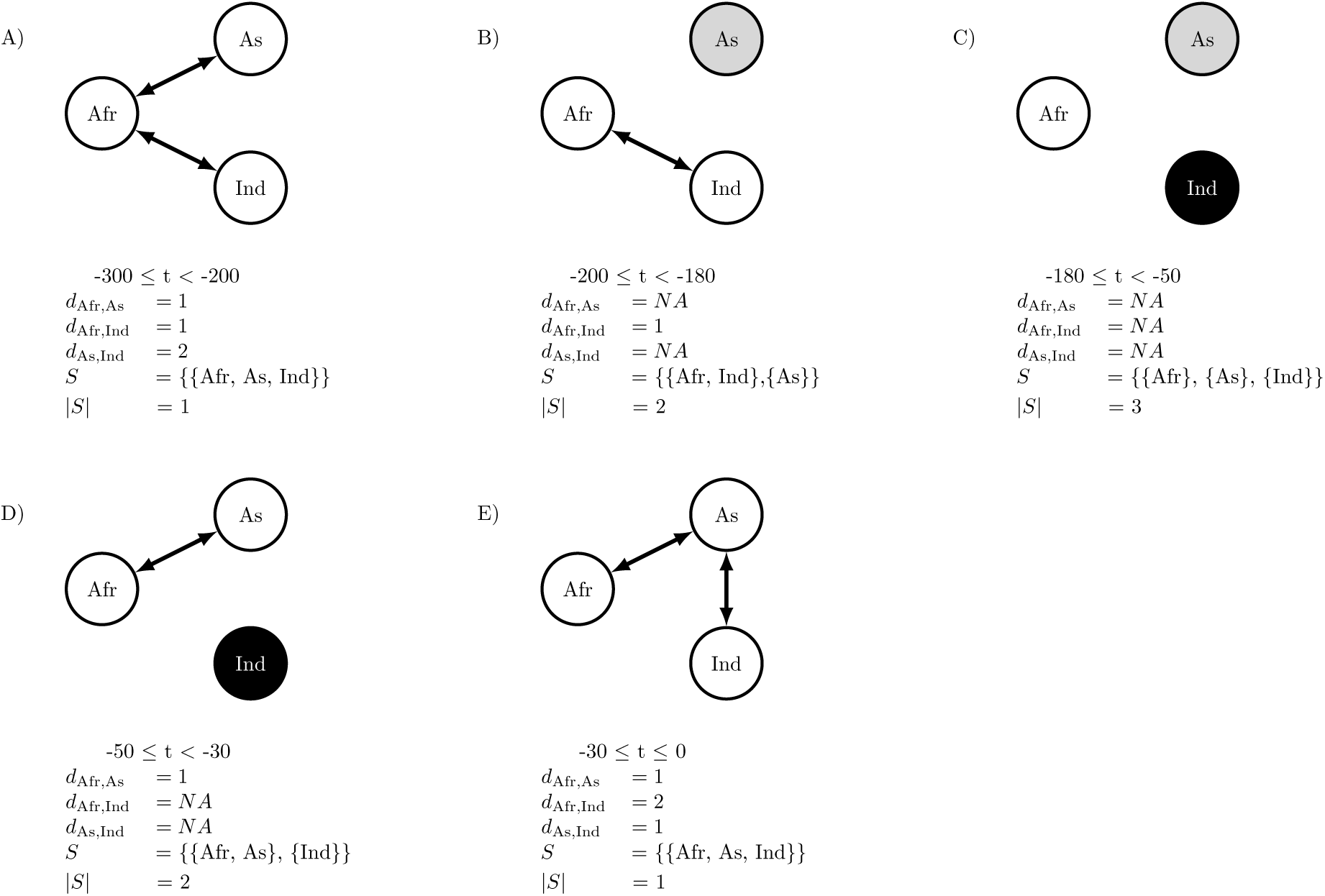
Biogeographic communicating classes. Dispersal routes shared by Africa (Afr), Asia (As), and India (Ind) are depicted for each time interval, *t*, over the past 300 Ma. Dispersal path lengths between areas *i* and *j* are given by *d_i,j_*, with NA meaning there is no route between areas (areas *i* and *j* are mutually unreachable). communicating classes per interval are given by *S* and by the shared coloring of areas (vertices), with |*S*| being the number of communicating classes.

Specifying communicating classes is partly difficult because we do not know the ease of dispersal between areas for most species throughout space and time. Encoding zero-valued dispersal rates directly into the model should be avoided given the apparent prevalence of long distance dispersal, sweepstakes dispersal, etc. across dispersal barriers (Carlquist 1966). Moreover, zero-valued rates imply that dispersal events between certain areas are not simply improbable but completely impossible, creating troughs of zero likelihood in the likelihood surface for certain dated-phylogeny-character patterns (Buerki et al. 2011). In a biogeographic dating framework, this might unintentionally eliminate large numbers of speciation scenarios from the space of possible hypotheses, resulting in distorted estimates. To avoid these problems, I take the dispersal graph as the weighted average of three distinct dispersal graphs assuming short, medium, or long distance dispersal modes, each with their own set of communicating classes (see Section 2.4).

Fundamentally, biogeographic dating depends on how rapidly a species may disperse between two areas, and how that dispersal rate changes over time. In one extreme case, dispersals between mutually unreachable areas do not occur after infinite time, and hence have zero probability. At the other extreme, when dispersal may occur between any pair of areas with equal probability over all time intervals, then paleogeography does not favor nor disfavor dispersal events (nor divergence events, implicitly) to occur during particular time intervals. In intermediate cases, so long as dispersal probabilities between areas vary across time intervals, the dispersal process informs when and what dispersal (and divergence) events occur. For instance, the transition probability of going from area *i* to *j* decreases as the average path length between *i* and *j* increases. During some time intervals, the average path length between two areas might be short, thus dispersal events occur more freely than when the average path is long. Comparing Figures 3A and 3E, the minimum number of events required to disperse from India to Africa is smaller during the Triassic (*t* = −250) than during the present (*t* = 0), and thus would have a relatively higher probability given the process operated for the same amount of time today (e.g. for a branch with the same length).

The concepts of adjacency, reachability, components, and communicating classes are not necessary to structure the rate matrix such that biogeographic events inform divergence times, though their simplicity is attractive. One could yield similar effects by parameterizing dispersal rates as functions of more complex area features, such as geographical distance between areas (Landis et al. 2013) or the size of areas (Tagliacollo et al. 2015). In this study, these concepts serve the practical purpose of summarizing perhaps the most salient feature of global paleogeography—that continents were not always configured as they were today—but also illuminate how time-heterogeneous dispersal rates produce transition probabilities that depend on geological time, which in turn inform the dates of speciation times.

### 2.3 Time-heterogeneous dispersal process

Let **Z** be a vector reporting biogeographic states for *M* > 2 taxa. The objective is to construct a time-heterogeneous CTMC where transition probabilities depend on time-calibrated paleogeographical features. For simplicity, species ranges are assumed to be endemic on the continental scale, so each taxon’s range may be encoded as an integer in *Z_i_* ∈ {1, 2,…, *N*}, where *N* is the number of areas.

The paleogeographical features that determine the dispersal process rates are assumed to be a piecewise-constant model, sometimes called a stratified (Ree et al. 2005; Ree and Smith 2008) or epoch model (Bielejec et al. 2014), where *K* − 1 breakpoints are dated in geological time to create *K* time intervals. These breakpoint times populate the vector, *τ* = (*τ*_0_ = −∞, *τ*_1_, *τ*_2_, …, *τ_K_*_–1_, *τ_K_* = 0), with the oldest interval spanning deep into the past, and the youngest interval spanning to the present.

While a lineage exists during the *k*^th^ time interval, its biogeographic characters evolve according to that interval’s rate matrix, **Q**(*k*), whose rates are informed by paleogeographical features present throughout time *τ_k_*_–1_ ≤ t < *τ_k_*. As a example of an paleogeographically-informed matrix structure, take **G**(*k*) to be a adjacency matrix indicating 1 when dispersal may occur between two areas and 0 otherwise, during time interval *k*. This adjacency matrix is equivalent to an undirected graph where areas are vertices and edges are dispersal routes. Full examples of **G** = (**G**(1), **G**(2),…, **G**(*K*)) describing Earth’s paleocontinental adjacencies are given in detail later.

With the paleogeographical vector **G**, I define the transition rates of **Q**(*k*) as equal to **G***_z_* (*k*). Similar rate matrices are constructed for all *K* time intervals that contain possible supported root ages for the phylogeny, Ψ. Figure 4 gives a simple example for three areas, where Asia shares positive dispersal rate with Africa when they are merged and no dispersal while split.

**Figure 4:**
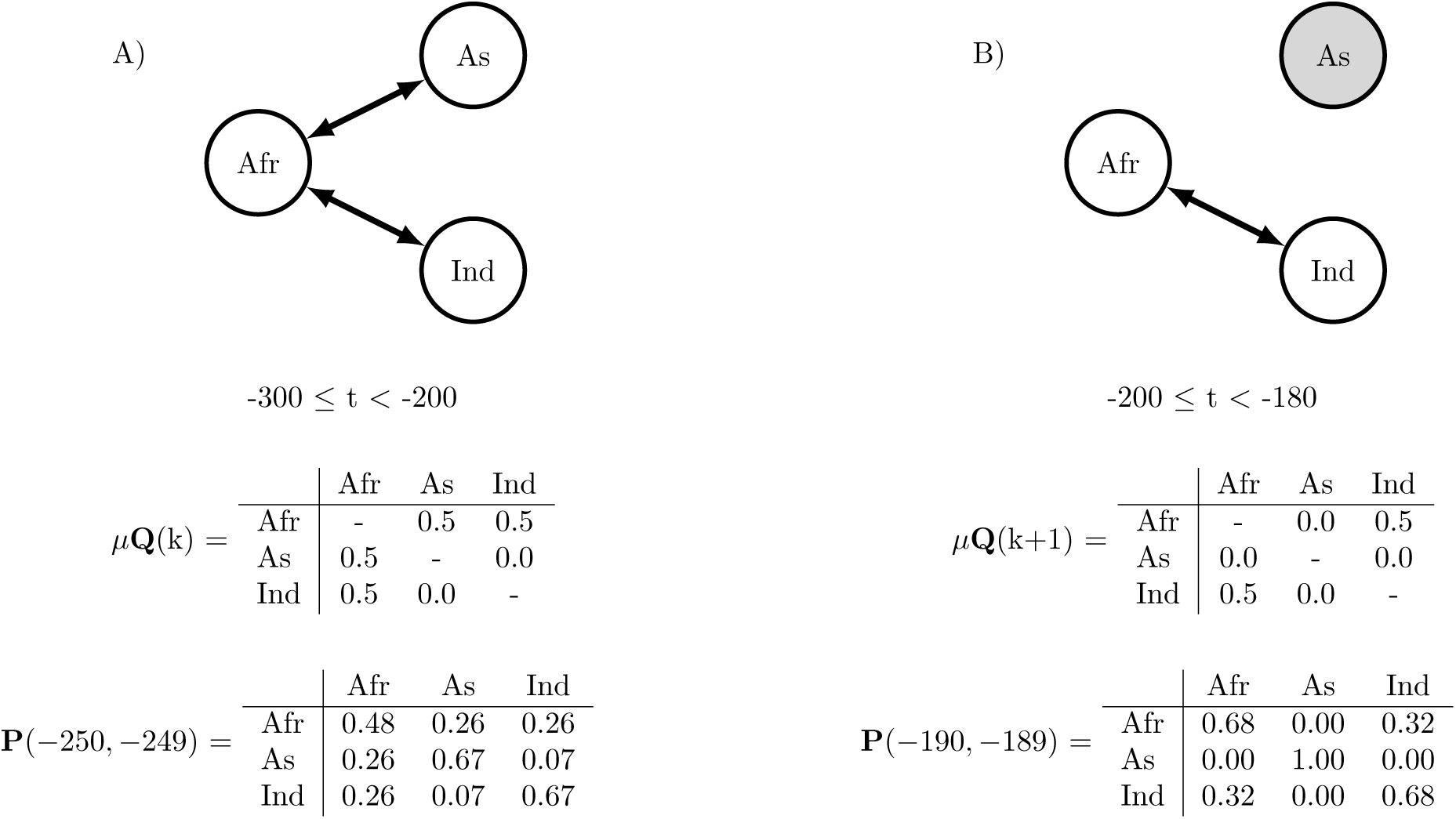
Piecewise-constant dispersal rate matrices. Dispersal routes shared by Africa (Afr), Asia (As), and India (Ind) are depicted for two time intervals, −300 ≤ t < −200 and −200 ≤ t < 180. Graphs and times correspond to those in A,B. Transition probabilities are computed for a unit time during different epochs with a time-homogeneous biogeographic clock rate *μ* = 0.5. A) The three areas are connected and all transitions have positive probability. B) As is unreachable from Afr and Ind, so transition probabilities into and out of As are zero.

For a piecewise-constant CTMC, the process’ transition probability matrix is the product of transition probability matrices spanning *m* breakpoints. To simplify notation, let *v* be the vector marking important times of events, beginning with the start time of the branch, *s*, followed by the *m* breakpoints satisfying *s* < *τ_k_* < *t*, ending the the end time of the branch, *t*, such that *v* = (*s*, *τ_k_*, *τ_k_*_+1_,…, *τ_k_*_+_*_m_*_–1_, *t*), and let *u*(*v_i_*, *τ*) be a “look-up” function that gives the index *k* that satisfies *τ_k_*_–1_ ≤ *v_i_* < *τ_k_*. The transition probabilty matrix over the intervals in v according to the piece-wise constant CTMC given by the vectors *τ* and **Q** is

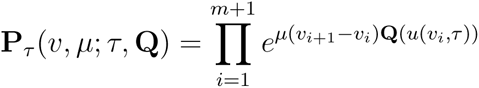

The pruning algorithm (Felsenstein 1981) is agnostic as to how the transition probabilties are computed per branch, so introducing the piecewise-constant CTMC does not prohibit the efficient computation of phylogenetic model likelihoods. See Bielejec et al. (2014) for an excellent review of piecewise-constant CTMCs as applied to phylogenetics.

In the above case, the times *s* and *t* are generally identifiable from *μ_z_* so long as **P***_τ_*(*v*, *μ*; *τ*, **Q**) ≠ **P***_τ_* (*v′*, *μ*′; *τ*, **Q**) for any supported values of *v*, *μ*, *v*′, and *μ*′. Note, I include *μ* as an explicit parameter in the transition probability matrix function for clarity, though they are suppressed in standard CTMC notation when *t* equals the product of rate and time, then the process effectively runs for the time, *μ*(*t* – *s*). For example, assume that **Q** is a time-homogeneous Jukes-Cantor model with no paleogeographical constraints, i.e. all transition rates are equal independent of *k*. The transition probability matrix for this model is readily computed via matrix exponentiation

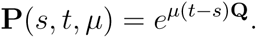

Note that **P**(*s*, *t*, *μ*) = **P***_τ_* (*v*, *μ*; *τ*, **Q**) when *v* = (*s*, *t*) – i.e. the time-heterogeneous process spans no breakpoints when *m* = 0 and is equivalent to a time-homogeneous process for the interval (*s*, *t*).

For a time-homogeneous model, multiplying the rate and dividing the branch length by the same factor results in an identical transition probability matrix. In practice this means the simple model provides no information for the absolute value of *μ* and the tree height of Ψ, since all branch rates could likewise be multiplied by some constant while branch lengths were divided by the same constant, i.e. **P**(*s*, *t*, 1) = **P**(*sμ*^−1^, *tμ*^−1^, *μ*). Similarly, since **P**(*s*, *t*, *μ*) = **P**(*s* + *c*, *t* + *c*, *μ*) for *c* ≥ 0, the absolute time when the process begins does not matter, only the amount of time that has elapsed. Extending a branch length by a factor of c requires modifying other local branch lengths in kind to satisfy time tree constraints, so the identifiability of the absolute time interval (*s*, *t*) depends on how “relaxed” (Drummond et al. 2006) the assumed clock and divergence time priors are with respect to the magnutude of *c*, which together induce some (often unanticipated) joint prior distribution on divergence times and branch rates (Heled and Drummond 2012; Warnock et al. 2015). In either case, rate and time estimates under the time-homogeneous process result from the induced prior distributions rather than by informing the process directly.

### 2.4 Adjacent-area terrestrial dispersal graph

I identified *K* = 26 times and *N* = 25 areas to capture the general features of continental drift and its effects on terrestrial dispersal (Figure 5; for all graphs and a link to the animation, see Supplemental Figure S1). All adjacencies were constructed visually, referencing Blakey (2006) and GPlates (Boyden et al. 2011), then corroborated using various paleogeographical sources (Table S2). The paleogeographical state per time interval is summarized as an undirected graph, where areas are vertices and dispersal routes are edges.

**Figure 5:**
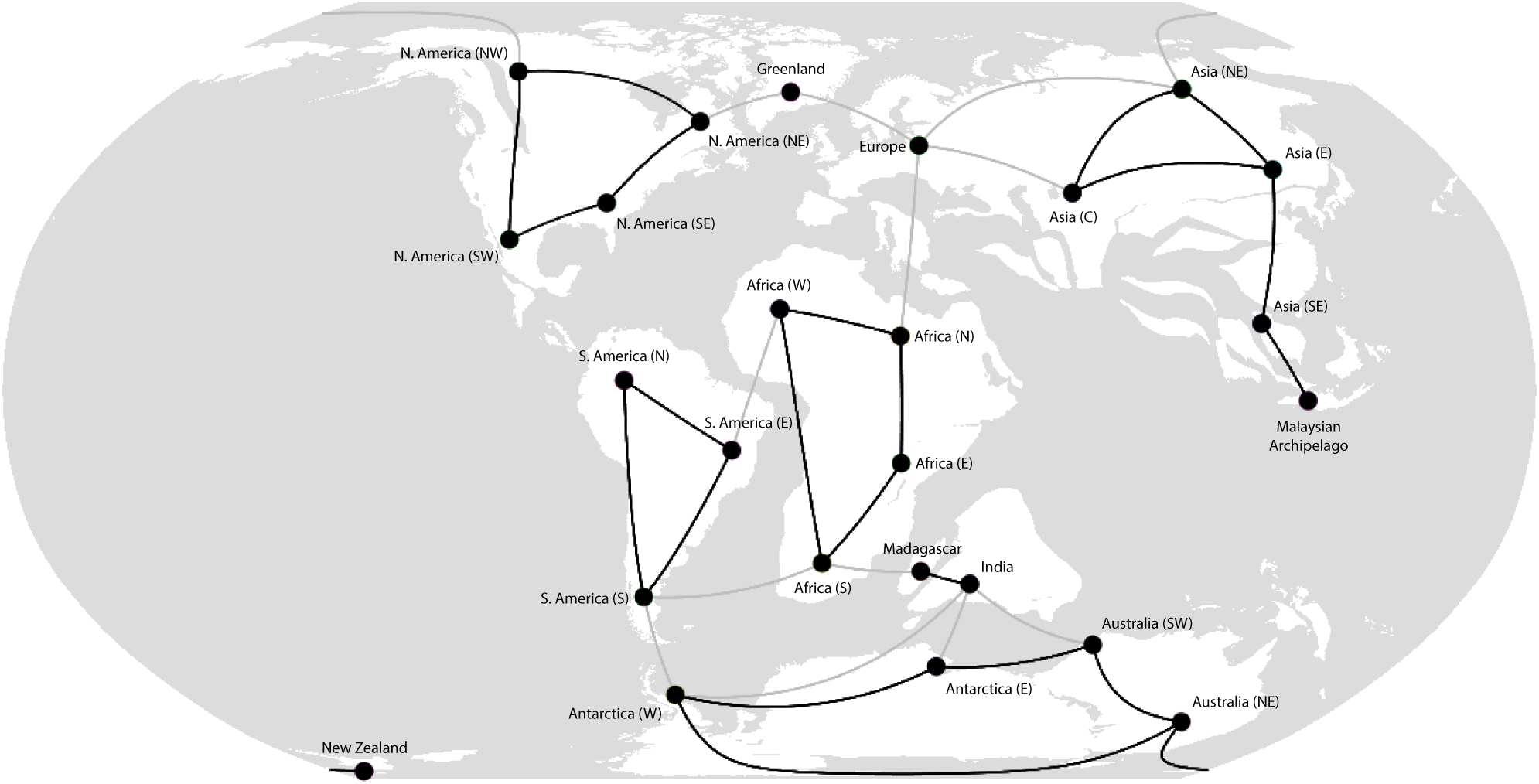
Dispersal graph for Epoch 14, 110–100Ma: India and Madagascar separate from Australia and Antarctica. A gplates (Gurnis et al. 2012) screenshot of Epoch 14 of 26 is displayed. Areas are marked by black vertices. Black edges indicate both short- and medium distance dispersal routes. Gray edges indicate exclusively medium distance dispersal routes. Long distance dispersal routes are not shown, but are implied to exist between all area-pairs. The short, medium, and long dispersal graphs have 8, 1, and 1 communicating classes, respectively. India and Madagascar each have only one short distance dispersal route, which they share. Both areas maintain medium distance dispersal routes with various Gondwanan continents during this epoch. The expansion of the Tethys Sea impedes dispersal into and out of Europe.

To proceed, I treat the paleogeographical states over time as a vector of adjacency matrices, where G_•_(*k*)*_i_,_j_* = 1 if areas *i* and *j* share an edge at time interval *k*, and G_•_(*k*)*_i_,_j_* = 0 otherwise. Temporarily, I suppress the time index, *k*, for the rate matrix Q(*k*), since all time intervals’ rate matrices are constructed in a similar manner. To mitigate the effects of model misspecification, **Q** is determined by a weighted average of three geological adjacency matrices

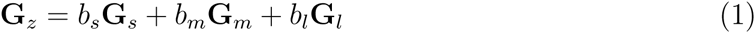

where *s*, *m*, and *l* correspond to short distance, medium distance, and long distance mode parameters.

Short, medium, and long distance dispersal processes encode strong, weak, and no geographical constraint, respectively. As distance-constrained mode weights *b_s_* and *b_m_* increase, the dispersal process grows more informative of the process’ previous state or communicating class (Figure 6). The vector of short distance dispersal graphs, **G***_s_* = (**G***_s_*(1), **G***_s_*(2),…, **G***_s_*(*K*)), marks adjacencies for pairs of areas allowing terrestrial dispersal without travelling through intermediate areas (Figure 6A). Medium distance dispersal graphs, **G***_m_*, include all adjacencies in **G***_s_* in addition to adjacencies for areas separated by lesser bodies of water, such as throughout the Malay Archipelago, while excluding transoceanic adjacencies, such as between South America and Africa (Figure 6B). Finally, long distance dispersal graphs, **G***_l_*, allow dispersal events to occur between any pair of areas, regardless of potential barrier (Figure 6C).

**Figure 6:**
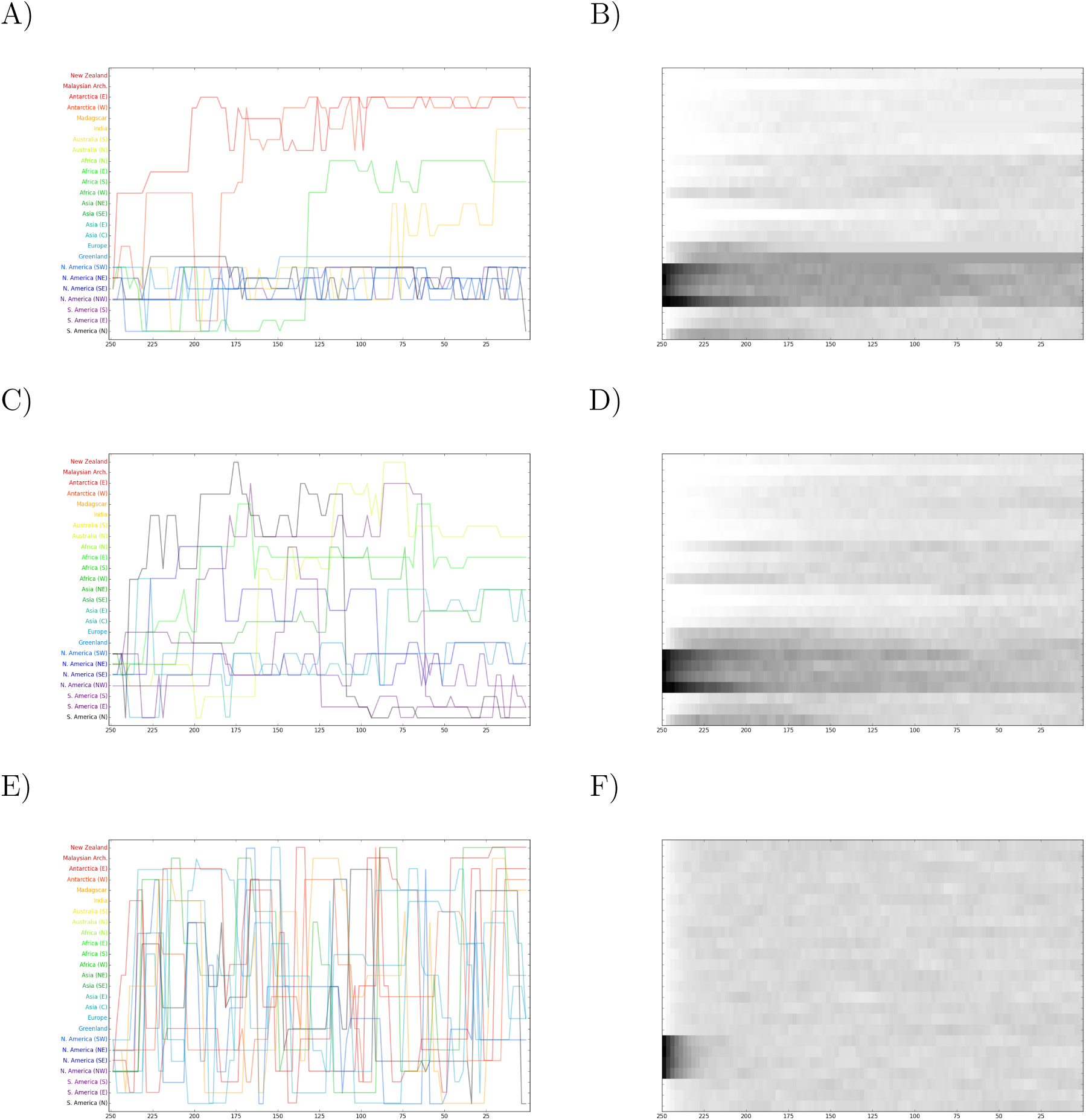
Sample paths for paleogeographically informed biogeographic process. The top, middle, and bottom panels show dispersal histories simulated by the pure short (A,B), medium (C,D), and long (E,F) distance process components. All processes originate in one of the four North American areas 250 Ma. The left column shows 10 of 2000 sample paths. Color indicates the area the lineage is found in the present (A,C,E). Colors for areas match those in Figure 8. The right column heatmap reports the sample frequencies for any of the 2000 dispersal process being in that state at that time (B,D,F).

To average over the three dispersal modes, *b_s_*, *b_m_*, and *b_l_* are constrained to sum to 1, causing all elements in **G***_z_* to take values from 0 to 1 (Eqn 1). Importantly, adjacencies specified by **G***_s_* always equal 1, since those adjacencies are also found in **G***_m_* and **G***_l_*. This means **Q** is a Jukes-Cantor rate matrix only when *b_l_* = 1, but becomes increasingly paleogeographically-structured as *b_l_* → 0. Non-diagonal elements of **Q** equal those of **G***_z_*, but are rescaled such that the average number of transitions per unit time equals 1, and diagonal elements of **Q** equal the negative sum of the remaining row elements. To compute transition probabilities, **Q** is later rescaled by a biogeographic clock rate, *μ*, prior to matrix exponentiation. The effects of the weights *b_s_*, *b_m_*, and *b_l_* on dispersal rates between areas are shown in Figure 7.

**Figure 7:**
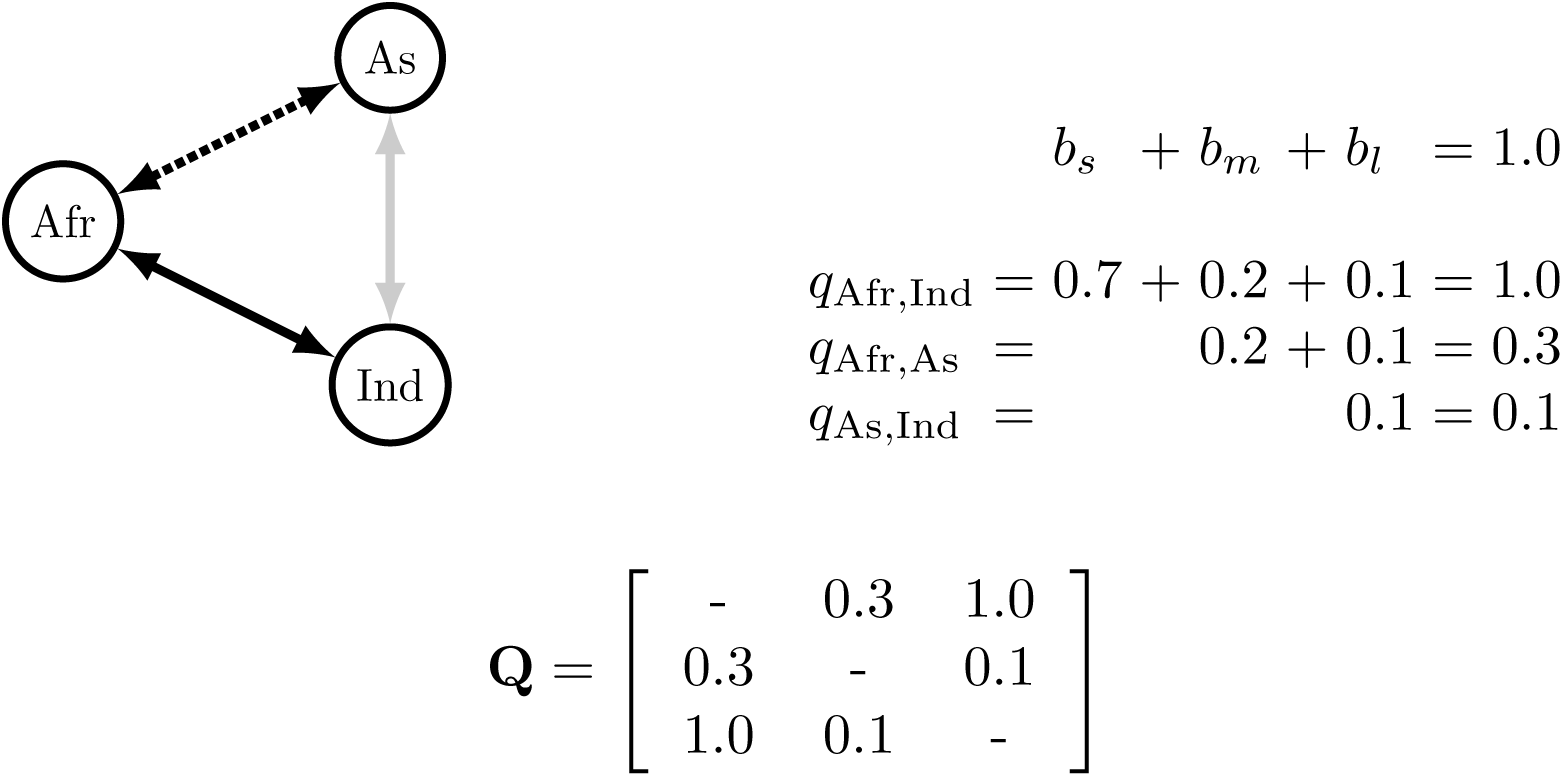
Example mode -weighted dispersal matrix. Short, medium, and long distance dispersal edges are represented by solid black, dashed black, and solid gray lines, respectively. Short, medium, and long distance dispersal weights are (*b_s_*, *b_m_*, *b_l_*) = (0.7, 0.2,0.1). The resulting mode-weighted dispersal matrix, **Q**, is computed with areas (states) ordered as (Afr, As, Ind). Afr and Ind share a short distance dispersal edge, therefore the dispersal weight is *b_s_* + *b_m_* + *b_l_* = 1.0. Afr and As share a medium distance edge with dispersal weight *b_m_* + *b_l_* = 0.3. Dispersal between As and Ind is only by long distance with weight *b_l_* = 0.1.

By the argument of that continental break-up (i.e. the creation of new communicating classes; Figure 1) introduces a bound on the minimum age of divergence, and that continental joining (i.e. unifying existing communicating classes; Figure 2) introduces a bound on the maximum age of divergence, then the paleogeographical model I constructed has the greatest potential to provide both upper and lower bounds on divergence times when the number of communicating classes is large, then small, then large again. This coincides with the formation of Pangaea, dropping from 8 to 3 communicating classes at 280 Ma, followed by the fragmentation of Pangaea, increasing from 3 to 11 communicating classes between 170 Ma and 100 Ma (Figure 8). It is important to consider this bottleneck in the number of communicating classes will be informative of root age only for fortuitous combinations of species range and species phylogeny. Just as some clades lack a fossil record, others are bound to lack a biogeographic record that is informative of origination times.

**Figure 8:**
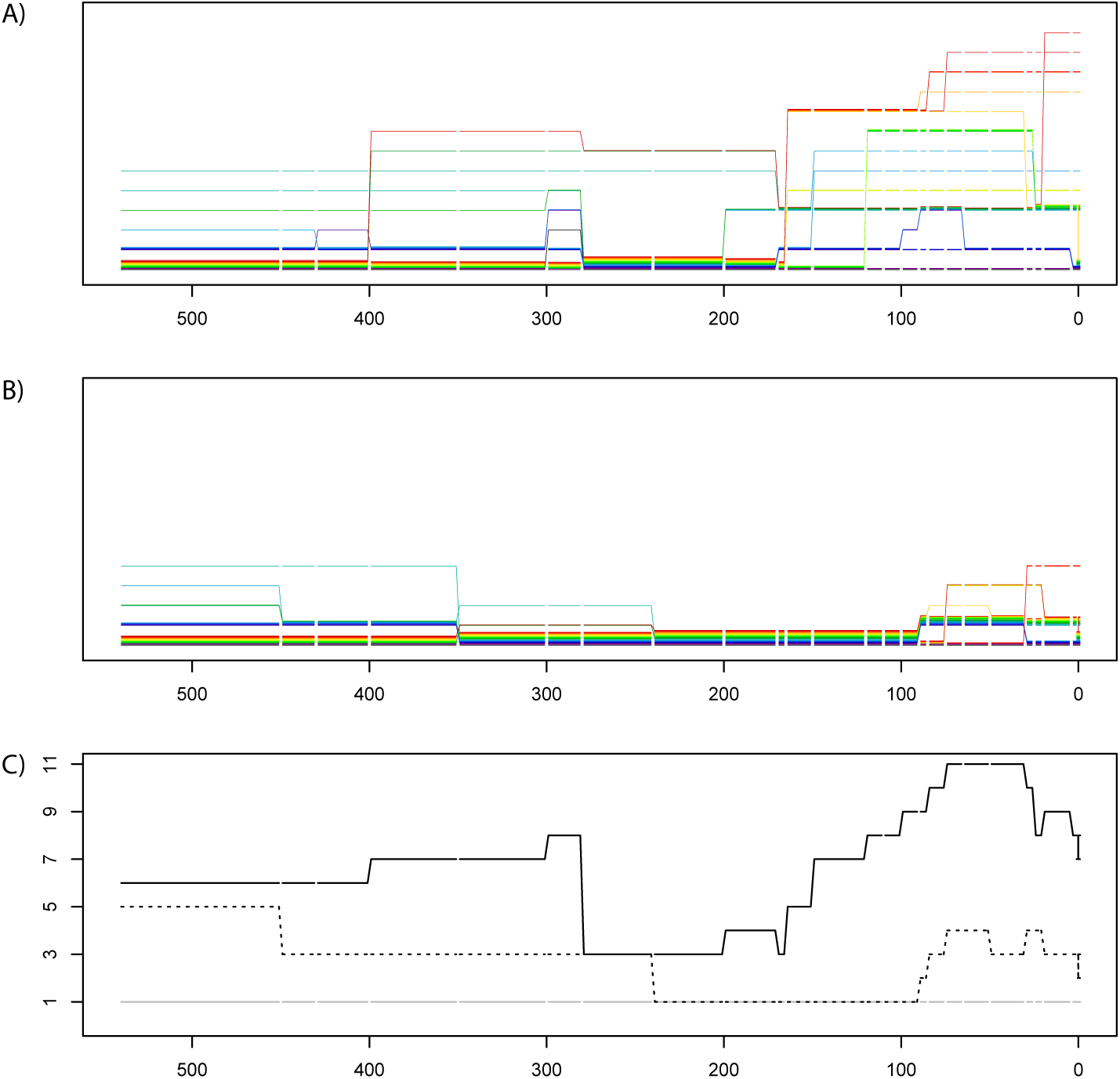
Dispersal graph properties summarized over time. Communicating classes of short distance dispersal graph (A) and medium distance dispersal graph (B) are shown. Each of 25 areas is represented by one line. Colors of areas match those listed in Figure 6. Grouped lines indicate areas in one communicating class during an interval of time. Vertical lines indicate transitions of areas joining or leaving communication clases, i.e. due to paleogeographical events. When no transition event occurs for an area entering a new epoch, the line is interrupted with gap. (C) Number of communicating classes: the black line corresponds to the short distance dispersal graph (A), the dotted line corresponds to medium distance dispersal graph (B), and the gray line corresponds to the long distance dispersal graph, which always has one communicating class.

## 3 Analysis

All posterior densities were estimated using Markov chain Monte Carlo (MCMC) as implemented in RevBayes, available at revbayes.com (Höhna et al. 2014). Data and analysis scripts are available at github.com/mlandis/biogeo_dating. Datasets are also available on Dryad at datadryad.org/XXX. Analyses were performed on the XSEDE supercomputing cluster (Towns et al. 2014).

## Simulation

Through simulation I tested whether biogeographic dating identifies rate from time. To do so, I designed the analysis so divergence times are informed solely from the molecular and biogeographic data and their underlying processes (Table 1). As a convention, I use the subscript *x* to refer to molecular parameters and z to refer to biogeographic parameters. Specifically, I defined the molecular clock rate as *μ_x_* = *e*/*r*, where e gives the expected number of molecular substitutions per site and *r* gives the tree height. Both *e* and *r* are distributed independently by uniform priors on (0; 1000). Biogeographic events occur with rate, 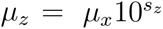 where *s_z_* has a uniform prior distribution on (−3; 3). To further subdue effects from the prior on posterior parameter estimates, the tree prior assigns equal probability to all node age distributions. No node calibrations were used. Each dataset was analyzed with (+G) and without (−G) the paleogeographic-dependent dispersal process.

**Table 1:**
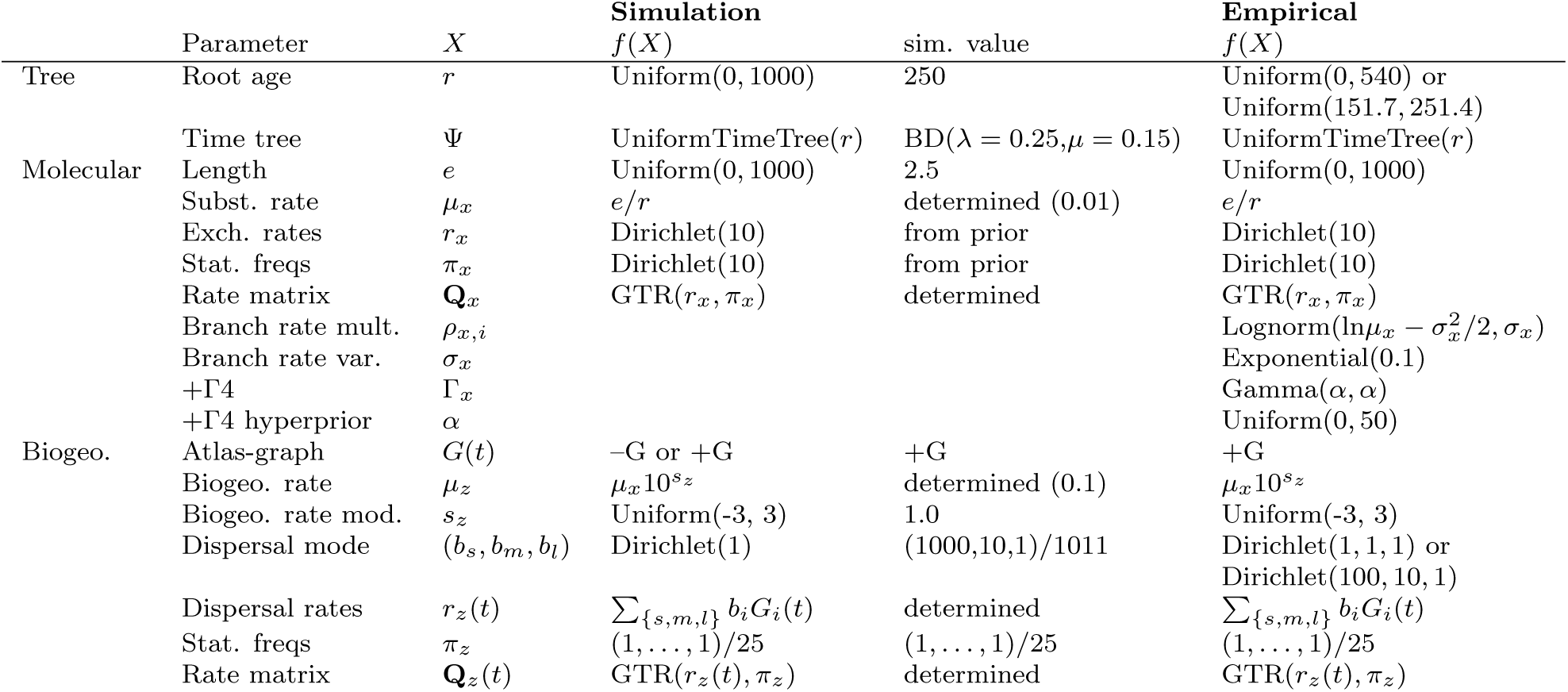
Model parameters. Model parameter names and prior distributions are described in the manuscript body. All empirical priors were identical to simulated priors unless otherwise stated. Priors used for the empirical analyses but not simulated analyses are left blank. Determined means the parameter value was determined by other model parameters.

Two further assumptions were made to simplify the analyses. First, although divergence times were free to vary, the tree topology was assumed to be known. Second, molecular and biogeographic characters evolve by strict global clocks. In principle, inferring the topology or using relaxed clock models should increase the variance in posterior divergence time estimates, but not greatly distort the performance of −G relative to +G.

Phylogenies with *M* = 50 extant taxa were simulated using a birth-death process with birth rate, *λ* = 0.25, and death rate, *μ* = 0.15, then rescaled so the root age equaled 250 Ma. Each dataset contained 500 nucleotides and 1 biogeographic character. Biogeographic characters were simulated under +G, where **G***_z_* is defined as piecewise-constant over 25 areas and 26 time intervals in the manner described in Section 2.4. In total, I simulated 100 datasets under the parameter values given in Table 1, where these values were chosen to reflect possible empirical estimates. Each dataset was analyzed under each of two models, then analyzed a second time to verify convergence (Gelman and Rubin 1992; Plummer et al. 2006). When summarizing posterior results, posterior mean-of-median and 95% highest posterior density (HPD95%) values were presented.

As expected, the results show the −G model extracts no information regarding the root age, so its posterior distribution equals its prior distribution, mean-of-median ≈ 499 (Figure 9A). In contrast, the +G model infers the mean-of-median root age 243 with a HPD95% interval width of 436, improving accuracy and precision in general.

**Figure 9:**
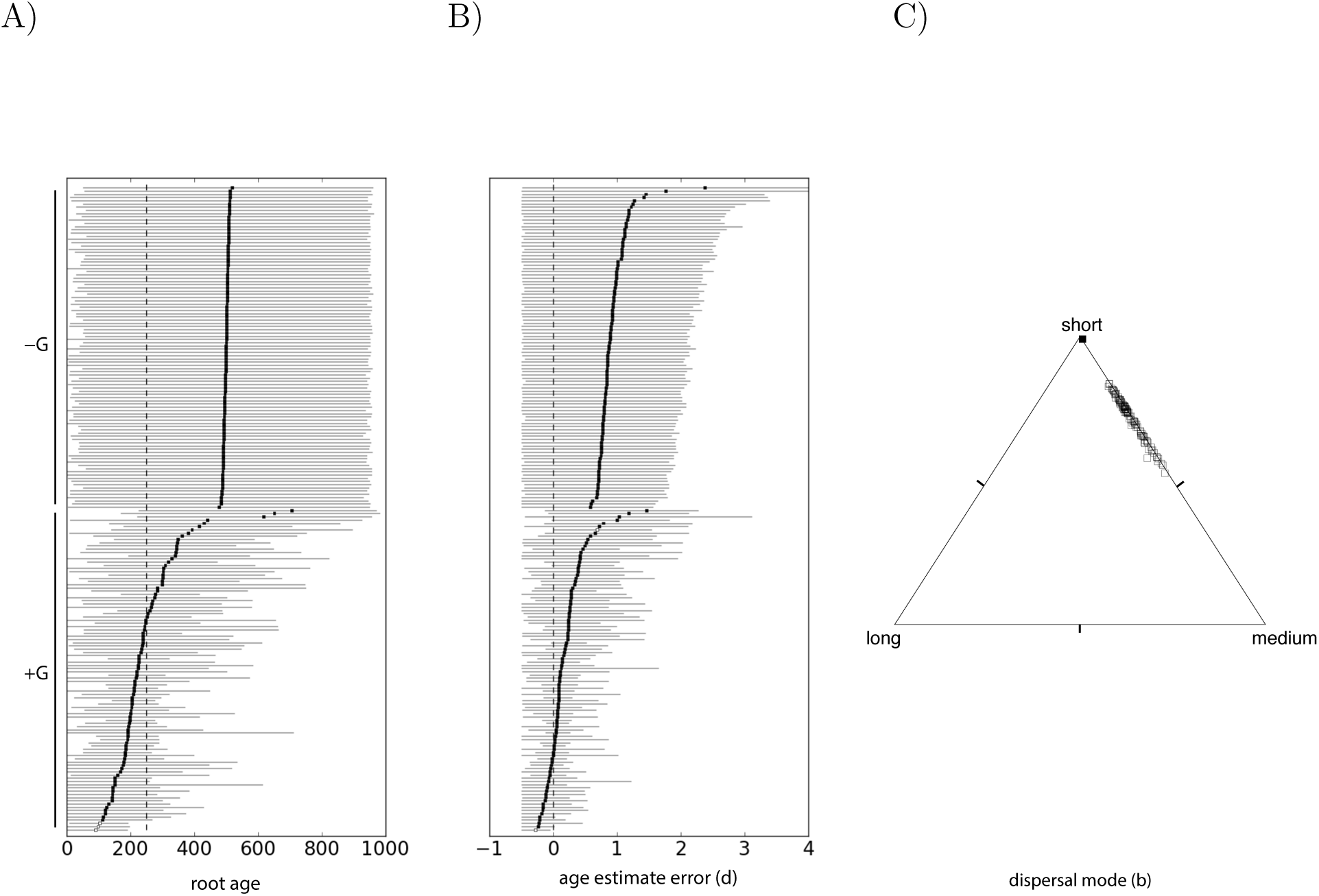
Posterior estimates for simulated data. A) Posterior estimates of root age. The true root age for all simulations is 250 Ma (dotted vertical line). B) Posterior estimates of relative node age error (Eqn 2). The true error term equals zero. Both A and B) −G analyses are on the top half, +G analyses are on the bottom. Each square marks the posterior mean root age estimate with the HPD95% credible interval. If the credible interval contains the true value, the square is filled. C) Posterior estimates of dispersal mode proportions for the +G simulations projected onto the unit 2-simplex. The filled circle gives the posterior median-of-medians, and the empty circles give posterior medians.

Estimated divergence time accuracy was assessed with the statistic

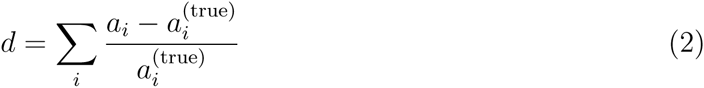

where *a* is a posterior sample of the node age vector and *a*_true_ is the true node age vector known through simulation. When *a* perfectly estimates *a*^(true)^ for all node ages, *d* = 0. When estimated node ages are too young (on average), *d* < 0, and when too old, *d* > 0. Inference under +G infers an mean *d* = 0.19 with a HPD95% interval width of ≈ 1.26, while −G performs substantially worse with *d* = 0.92 and width ≈ 2.75 (Figure 9B). Posterior estimates generally favored short over medium and long distance dispersal as was assumed under simulation (Figure 9C). Dispersal mode parameter estimates were (*b_s_*, *b_m_ b_l_*) = (0.766, 0.229, 0.003), respectively, summarized as median-of-medians across simulated replicates.

## **Empirical:** *Testudines*

To assess the accuracy of the method, I performed a biogeographic dating analysis on extant turtle species (*Testudines*). Extant turtles fall into two clades, *Pleurodira,* found in the Southern hemisphere, and *Cryptodira,* found predominantly in the Northern hemisphere. Their modern distribution shadows their biogeographic history, where *Testudines* are thought to be Gondwanan in origin with the ancestor to cryptodires dispersing into Laurasia during the Jurassic (Crawford et al. 2015). Since turtles preserve so readily in the fossil record, estimates of their phylogeny and divergence times have been profitably analyzed and reanalyzed by various researchers (Joyce 2007; Hugall et al. 2007; Danilov and Parham 2008; Alfaro et al. 2009; Dornburg et al. 2011; Joyce et al. 2013; Sterli et al. 2013; Warnock et al. 2015). This makes them ideal to assess the efficacy of biogeographic dating, which makes no use of their replete fossil record: if both biogeography-based and fossil-based methods generate similar results, they co-validate each others’ correctness (assuming they are not both biased in the same manner).

To proceed, I assembled a moderately sized dataset. First, I aligned cytochrome B sequences for 185 turtle species (155 cryptodires, 30 pleurodires) using MUSCLE 3.8.31 (Edgar 2004) under the default settings. Assuming the 25-area model presented in Section 2.4, I consulted GBIF (gbif.org) and IUCN Red List (iucnredlist.org) to record the area(s) in which each species was found. Species occupying multiple areas were assigned ambiguous tip states for those areas. Missing data entries were assigned to the six sea turtle species used in this study to effectively eliminate their influence on the (terrestrial) biogeographic process. To simplify the analysis, I assumed the species tree topology was fixed according to Guillon et al. (2012), which was chosen for species coverage, pruning away unused taxa. All speciation times were considered random variables to be estimated. The tree topology and biogeographic states are shown in Supplemental Figure S2. All data are recorded on datadryad.org/XXX.

Like the simulation study, my aim is to show that the paleogeographically-aware +G model identifies the root age in units of absolute time. To reiterate, the posterior root age should be identical to the prior root age when the model cannot inform the root age. If the prior and posterior differ, then the data under the model are informative. The root age was constrained to Uniform(0, 540), forbidding the existence of Precambrian turtles. To improve biological realism, I further relaxed assumptions about rate variability for the molecular model of evolution, both among sites (Yang 1994) and across branches (Lepage et al. 2007; Drummond et al. 2006) (Table 1).

Biogeographic dating infers a posterior median root age of 201 with HPD95% credible interval of (115, 339) (Figure 10A). This is consistent current root age estimates informed from the fossil record (Figure 11). The posterior mode of dispersal mode is (*b_s_, b_m_, b_l_*) = (0.47, 0.51, 0.02), with short and medium distance dispersal occurring at approximately equal rates and long distance dispersal being rare by comparison. Biogeographic events occurred at a ratio of about 6:1 when compared to molecular events (posterior means: *μ_x_* = 1.9E−3, *μ_z_* = 1.1E−2). The posterior mode tree height measured in expected number of dispersal events is 2.3 with HPD95% (1.5, 3.0), i.e. as a treewide average, the current location of each taxon is the result of about two dispersal events.

**Figure 10:**
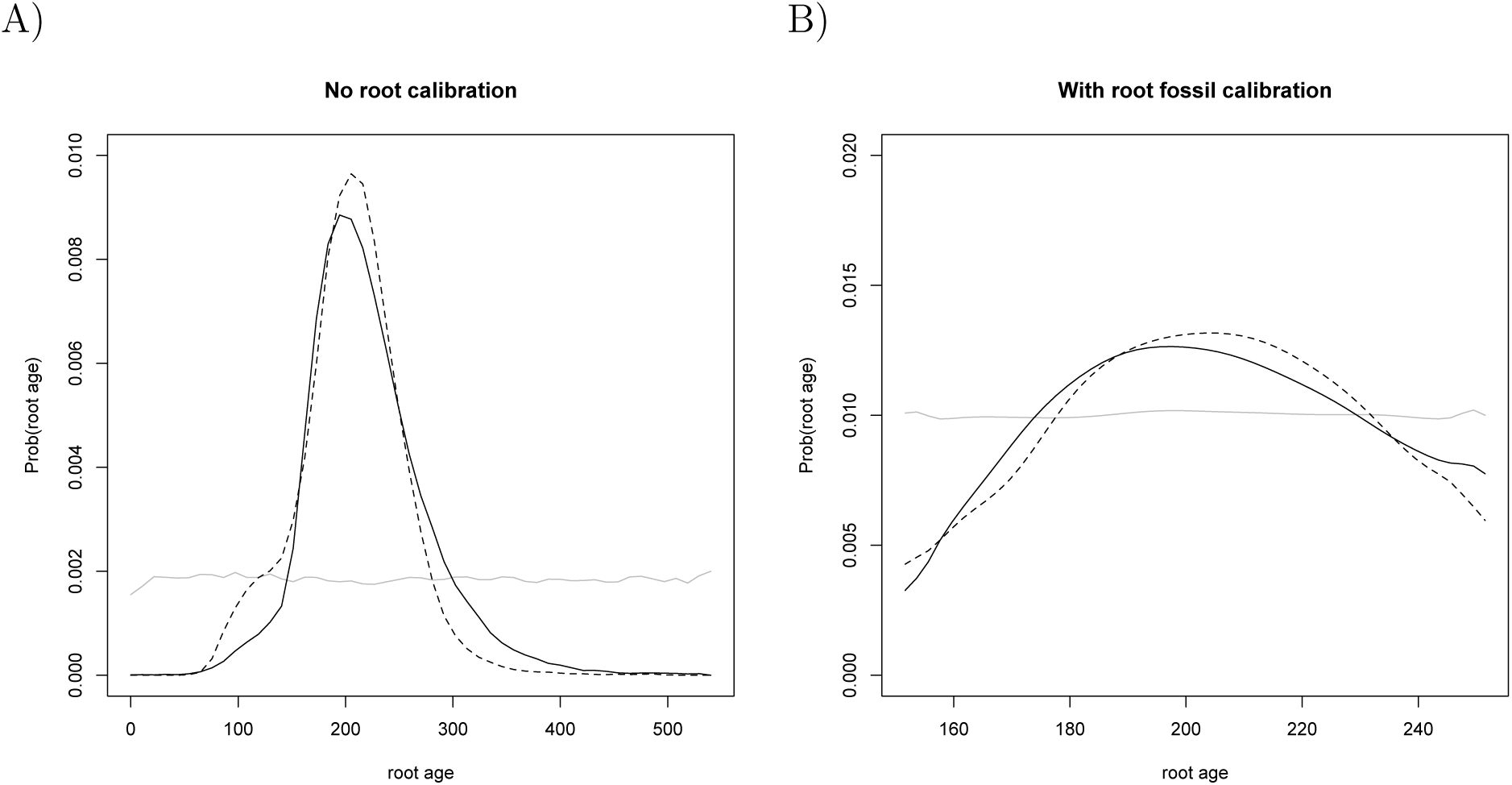
Posterior root age of turtles by biogeographic dating. Six root age posterior estimates were computed using biogeographic dating, each using variations on flat or short-biased priors for key parameters. Figure A assumes no knowledge of fossils with Uniform(0, 540) root age prior. Figure B follows Joyce et al. (2013) and assumes Uniform(151.7, 251.4) as a root node age calibration. The black solid posterior density assumes a flat prior on dispersal mode. The black dotted posterior density assumes an short-biased prior Dirichlet(100, 10, 1) on dispersal mode. The gray solid posterior density ignores paleogeography.

**Figure 11:**
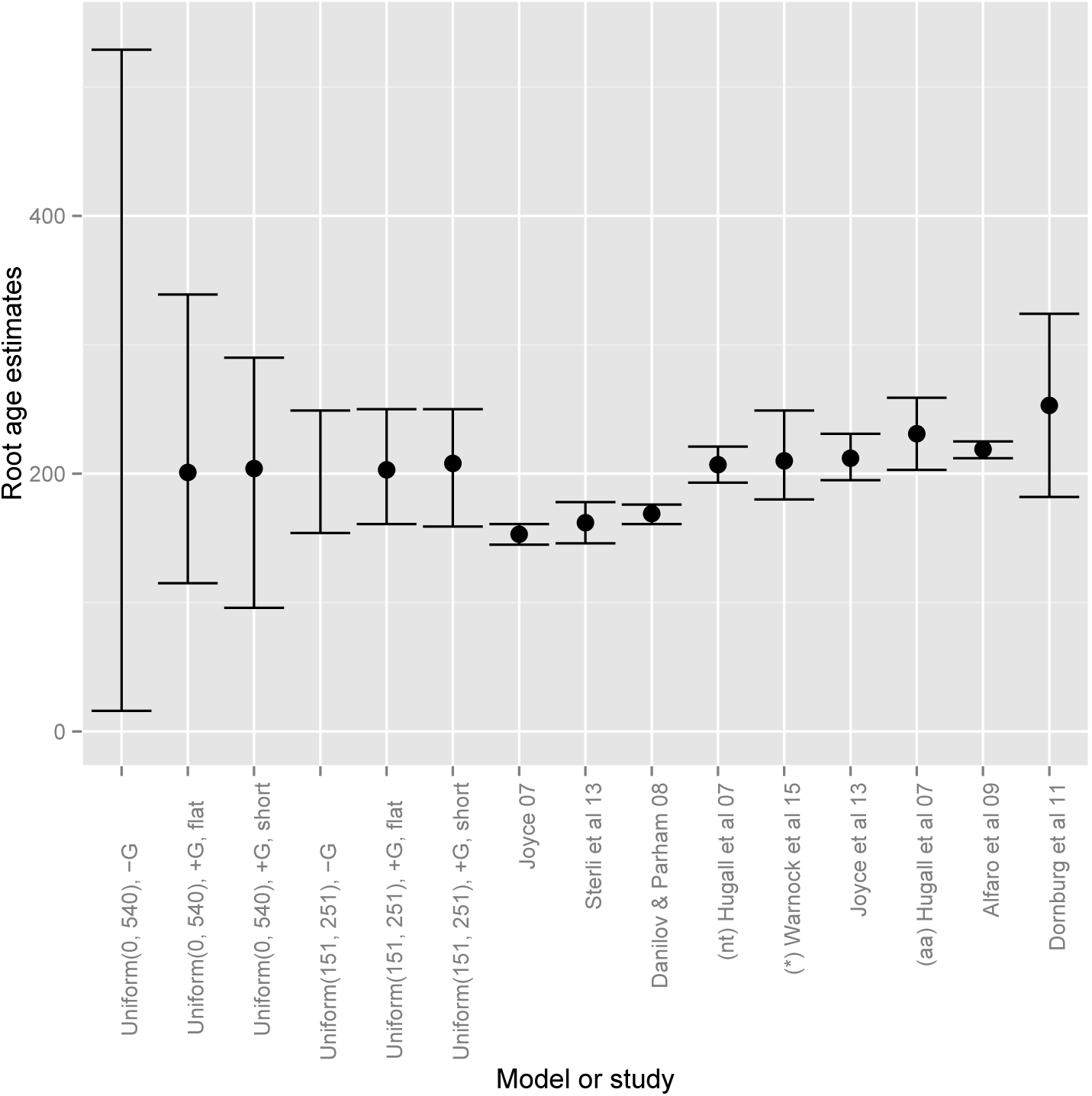
Root age comparison. Root age estimates are presented both for analysis conducted for this manuscript and as reported in existing publications. Existing estimates are as reported in Sterli et al. (2013) and supplemented recently reported results. Points and whiskers correspond to the point estimates and estimate confidence, which varies across analyses. The six left estimates were computed using biogeographic dating, each using variations on flat or short-biased priors for key parameters. Two of these analyses ignore paleogeography (−G) so the posterior root age is the uniform prior root age, whose mode (not shown) equals all values supported by the prior. Hugall et al. (2007) reports ages for analyses using amino acids (aa) and nucleotides (nt). Warnock et al. (2015) reports many estimates while exploring prior sensitivity, but only uniform prior results are shown here.

The flat prior distribution for competing dispersal modes is Dirichlet(1, 1,1) and does not capture the intuition that short distance dispersal should be far more common than long distance dispersal. I encoded this intuition in the dispersal mode prior, setting the distribution to Dirichlet(100, 10, 1), which induces expected proportion of 100:10:1 short:medium:long dispersal events. After re-analyzing the data with the short-biased dispersal prior, the posterior median and HPD95% credible interval were estimated to be, respectively, 204 (96, 290) (Figure 10A).

Biogeographic dating is compatible with fossil dating methods, so I repeated the analysis for both flat and informative prior dispersal modes while substituting the Uniform(0, 540) prior on root age calibration for Uniform(151.7, 251.4) (Joyce et al. 2013). When taking biogeography into account, the model more strongly disfavors post-Pangaean origins for the clade than when biogeography is ignored, but the effect is mild. Posterior distributions of root age was relatively insensitive to the flat and short-biased dispersal mode priors, with posterior medians and credible intervals of 203 (161, 250) and 208 (159, 250), respectively.

All posterior root state estimates favored South America (N) for the paleogeographically-informed analyses (Figure 12A). Although this is in accord with the root node calibration adopted from Joyce et al. (2013)—*Caribemys oxfordiensis,* sampled from Cuba, and the oldest accepted crown group testudine—the fossil is described as a marine turtle, so the accordance may simply be coincidence. In contrast, the paleogeographically-naive models support Southeast Asian origin of *Testudines*, where, incidentally, Southeast Asia is the most frequently inhabited area among the 185 testudines. For the analysis with a flat dispersal mode prior and no root age calibration, all root states with high posterior probability appear to concur on the posterior root age density (Figure 12B), i.e. regardless of conditioning on South America (N), North America (SE), or North America (SW) as a root state, the posterior root age density is roughly equal.

**Figure 12:**
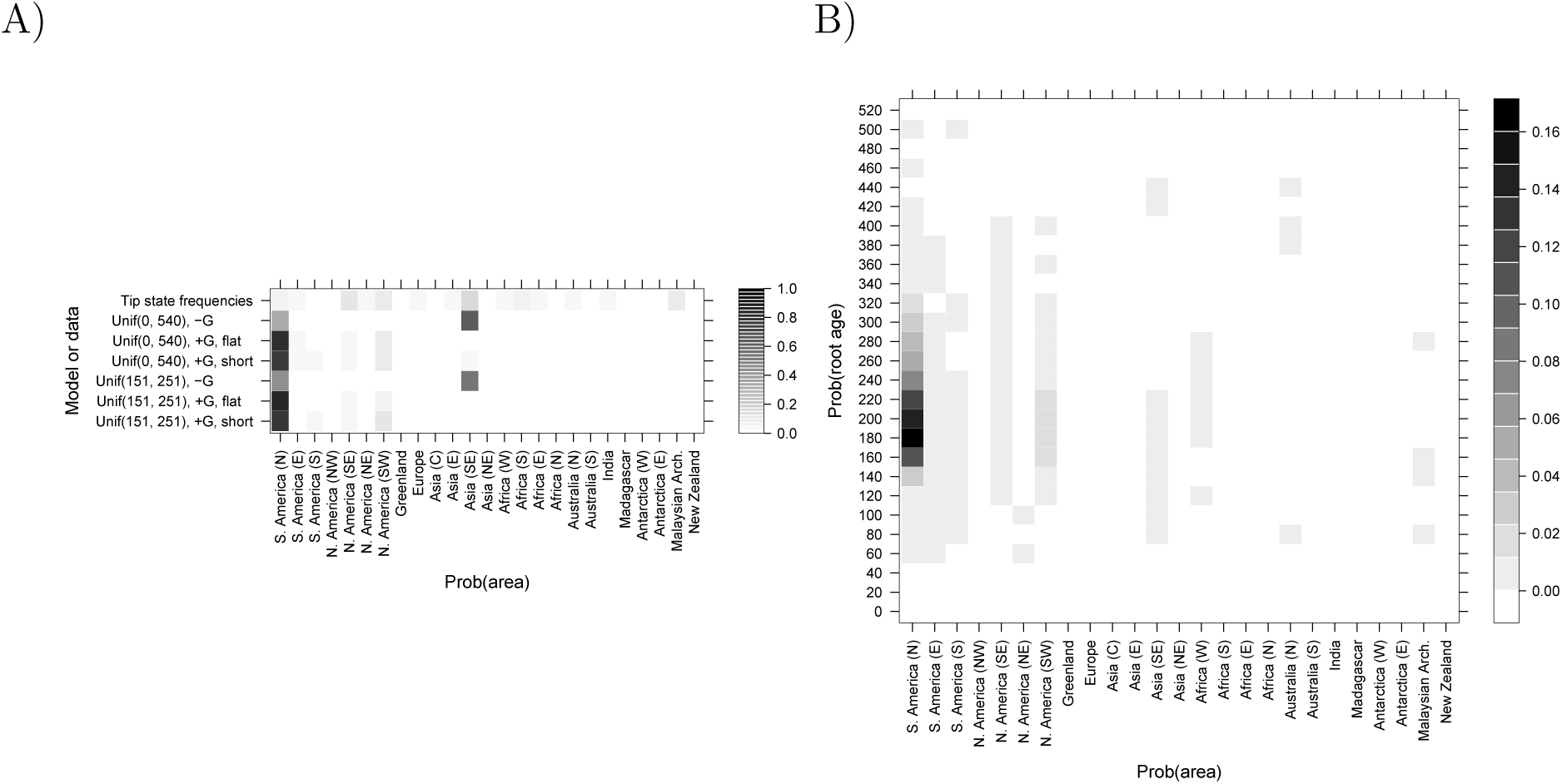
Root state estimates. A) Posterior probabilities of root state are given for the six empirical analyses. B) Joint-marginal posterior probabilities of root age and root state are given for the empirical analysis without a root calibration and with a flat dispersal mode prior. Root ages are binned into intervals of width 20.

## 4 Discussion

The major obstacle preventing the probabilistic union of paleogeographical knowledge, biogeographic inference, and divergence time estimation has been methodological, which I have attempted to redress in this manuscript. The intuition justifying prior-based fossil calibrations (Parham et al. 2011), i.e. that fossil occurrences should somehow inform divergence times, has recently been formalized into several models (Pyron 2011; Ronquist et al. 2012; Heath et al. 2014). Here I present an analogous treatment for prior-based biogeographic calibrations, i.e. that biogeographic patterns of modern species echo time-calibrated paleobiogeographic events, by describing how epoch models (Ree et al. 2005; Ree and Smith 2008; Bielejec et al. 2014) are informative of absolute divergence times. Briefly, I accomplished this using a simple time-heterogeneous dispersal process (Sanmartín et al. 2008), where dispersal rates are piecewise-constant, and determined by a graph-based paleogeographical model (Section 2.4). The paleogeographical model itself was constructed by translating various published paleogeographical reconstructions (Figure 5) into a time-calibrated vector of dispersal graphs.

Through simulation, I showed biogeographic dating identifies tree height from the rates of molecular and biogeographic character change. This simulation framework could easily be extended to investigate for what phylogenetic, paleogeographic, and biogeographic conditions one is able to reliably extract information for the root age. For example, a clade with taxa invariant for some biogeographic state would contain little to no information about root age, provided the area has always existed and had a constant number of dispersal edges over time. At the other extreme, a clade with a very high dispersal rate or with a proclivity towards long distance dispersal might provide little due to signal saturation (Figure 6C). The breadth of applicability of biogeographic dating will depend critically on such factors, but because we do not expect to see closely related species uniformly distributed about Earth nor in complete sympatry, that breadth may not be so narrow, especially in comparison to the fossil record.

The majority of groups have poor fossil records, and biogeographic dating provides a second hope for dating divergence times. Since biogeographic dating does not rely on any fossilization process or data directly, it is readily compatible with existing fossil-based dating methods (Figure 10B). When fossils with geographical information are available, researchers have shown fossil taxa improve biogeographical analyses (Moore et al. 2008; Wood et al. 2012). In principle, the biogeographic process should guide placement of fossils on the phylogeny, and the age of the fossils should improve the certainty in estimates of ancestral biogeographic states (Slater et al. 2012), on which biogeographic dating relies. Joint inference of divergence times, biogeography, and fossilization stands to resolve recent paleobiogeographic conundrums that may arise when considering inferences separately (Beaulieu et al. 2013; Wilf and Escapa 2014).

Because time calibration through biogeographic inferences comes primarily from the paleogeographical record, not the fossil record, divergence times may be estimated from exclusively extant taxa under certain biogeographical and phylogenetic conditions. When fossils are available, however, biogeographic dating is compatible with other fossil-based dating methods (e.g. node calibrations, fossil tip dating, fossilized birth-death). As a proof of concept, I assumed a flat root age calibration prior for the origin time of turtles: the posterior root age was also flat when paleogeography was ignored, but Pangaean times of origin were strongly preferred when dispersal rates conditioned on paleogeography (Figure 10). Under the uninformative prior distributions on root age, biogeographic dating estimated turtles originated between the Mississipian (339 Ma) and Early Cretaceous (115 Ma) periods, with a median age of 201 Ma. Under an ignorance prior where short, medium, and long distance dispersal events have equal prior rates, short and medium distance dispersal modes are strongly favored over long distance dispersal. Posterior estimates changed little by informing the prior to strongly prefer short distance dispersal. Both with and without root age calibrations, and with flat and biased dispersal mode priors, biogeographic dating placed the posterior mode origin time of turtles at approximately 210-200 Ma, which is consistent with fossil-based estimates (Figure 11).

### Model inadequacies and future extensions

The simulated and empirical studies demonstrate biogeographic dating improves divergence time estimates, with and without fossil calibrations, but many shortcomings in the model remain to be addressed. When any model is misspecified, inference is expected to produce uncertain, or worse, spurious results (Lemmon and Moriarty 2004), and biogeographic models are not exempted. I discuss some of the most apparent model misspecifications below.

Anagenetic range evolution models that properly allow species inhabit multiple areas should improve the informativeness of biogeographic data. Imagine taxa *T*1 and *T*2 inhabit areas *ABCDE* and *FGHIJ*, respectively. Under the simple model assumed in this paper, the tip states are ambiguous with respect to their ranges, and for each ambiguous state only a single dispersal event is needed to reconcile their ranges. Under a pure anagenetic range evolution model (Ree et al. 2005), at least five dispersal events are needed for reconciliation. Additionally, some extant taxon ranges may span ancient barriers, such as a terrestrial species spanning both north and south of the Isthmus of Panama. This situation almost certainly requires a dispersal event to have occurred after the isthmus was formed when multiple-area ranges are used. For single-area species ranges coded as ambiguous states, the model is incapable of evaluating the likelihood that the species is found in both areas simultaneously, so additional information about the effects of the paleogeographical event on divergence times is potentially lost.

Any model where the diversification process and paleogeographical states (and events) are correlated will obviously improve divergence time estimates so long as that relationship is biogeographically realistic. Although the repertoire of cladogenetic models is expanding in terms of types of transition events, they do not yet account for geographical features, such as continental adjacency or geographical distance. Incorporating paleogeographical structure into cladogenetic models of geographically-isolated speciation, such as vicariance (Ronquist 1997), allopatric speciation (Ree et al. 2005; Goldberg et al. 2011), and jump dispersal (Matzke 2014), is crucial not only to generate information for biogeographic dating analyses, but also to improve the accuracy of ancestral range estimates. Ultimately, cladogenetic events are state-dependent speciation events, so the desired process would model range evolution jointly with the birth-death process (Maddison et al. 2007; Goldberg et al. 2011), but inference under these models for large state spaces is currently infeasible. Regardless, any cladogenetic range-division event requires a widespread range, which in turn implies it was preceeded by dispersal (range expansion) events. Thus, if we accept that paleogeography constrains the dispersal process, even a simple dispersal-only model will extract dating information when describing a far more complex evolutionary process.

That said, the simple paleogeographical model described herein (Section 2.4) has many shortcomings itself. It is only designed for terrestrial species originating in the last 540 Ma. Rates of dispersal between areas are classified into short, medium, and long distances, but with subjective criteria. The number of epochs and areas was limited by my ability to comb the literature for well-supported paleogeological events, while constrained by computational considerations. The timing of events was assumed to be known perfectly, despite the literature reporting ranges of estimates. Certainly factors such as global temperature, precipitation, ecoregion type, etc. affect dispersal rates between areas, but were ignored. All of these factors can and should be handled more rigorously in future studies by modeling these uncertainties as part of a joint Bayesian analysis (Höhna et al. 2014).

Despite these flaws, defining the paleogeographical model serves as an excercise to identify what features allow a biogeographic process to inform speciation times. Dispersal barriers are clearly clade-dependent, e.g. benthic marine species dispersal would be poorly modeled by the terrestrial graph. Since dispersal routes for the terrestrial graph might serve as dispersal barriers for a marine graph, there is potential for learning about mutually exclusive dispersal corridor use in a multi-clade analysis (Sanmartín et al. 2008). Classifying dispersal edges into dispersal mode classes may be made rigorous using clustering algorithms informed by paleogeographical features, or even abandoned in favor of modeling rates directly as functions of paleogeographical features like distance. Identifying significant areas and epochs remains challenging, where presumably more areas and epochs are better to approximate continuous space and time, but this is not without computational challenges (Ree and Sanmartín 2009; Webb and Ree 2012; Landis et al. 2013). Rather than fixing epoch event times to point estimates, one might assign empirical prior distributions based on collected estimates. Ideally, paleogeographical event times and features would be estimated jointly with phylogenetic evidence, which would require interfacing phylogenetic inference with paleogeographical inference. This would be a profitable, but substantial, interdisciplinary undertaking.

## Conclusion

Historical biogeography is undergoing a probabilistic renaissance, owing to the abundance of georeferenced biodiversity data now hosted online and the explosion of newly published biogeographic models and methods (Ree et al. 2005; Ree and Smith 2008; Sanmartín et al. 2008; Lemmon and Lemmon 2008; Lemey et al. 2010; Goldberg et al. 2011; Webb and Ree 2012; Landis et al. 2013; Matzke 2014; Tagliacollo et al. 2015). Making use of these advances, I have shown how patterns latent in biogeographic characters, when viewed with a paleogeographic perspective, provide information about the geological timing of speciation events. The method conditions directly on biogeographic observations to induce dated node age distributions, rather than imposing (potentially incorrect) beliefs about speciation times using node calibration densities, which are data-independent prior densities. Biogeographic dating may present new opportunities for dating phylogenies for fossil-poor clades since the technique requires no fossils. This establishes that historical biogeography has untapped practical use for statistical phylogenetic inference, and should not be considered of secondary interest, only to be analysed after the species tree is estimated.

## Acknowledgements

I thank Tracy Heath, Sebastian Höhna, Josh Schraiber, Lucy Chang, Pat Holroyd, Nick Matzke, Michael Donoghue, and John Huelsenbeck for valuable feedback, support, and encouragement regarding this work. I also thank James Albert, Alexandre Antonelli, and the Society of Systematic Biologists for inviting me to present this research at Evolution 2015 in Guarujá, Brazil. Simulation analyses were computed using XSEDE, which is supported by National Science Foundation grant number ACI-1053575.

